# Identification of regulatory promoter sequences directing *MtCP6* transcription at the onset of nodule senescence *in Medicago truncatula*

**DOI:** 10.1101/2024.11.25.625165

**Authors:** Li Yang, Lisa Frances, Fernanda de Carvalho-Niebel, Pierre Frendo, Eric Boncompagni

## Abstract

- The symbiotic association of legumes with rhizobia results in the formation of new root organs called nodules. However, the lifespan of nodules is limited by the senescence process. Increased proteolytic activity is one of the hallmarks of nodule senescence. In *Medicago truncatula*, a papain cysteine protease encoding gene, *MtCP6*, is a marker for the onset of nodule senescence under both developmental and stress-induced pathways.
- To identify promoter regions conferring *MtCP6* senescence-related expression, progressive *MtCP6* promoter deletions were generated and the resulting sequences were fused with a reporter gene for promoter::GUS fusion analysis in transgenic *M. truncatula* roots.
- *In planta,* a minimal promoter sequence of 67 bp was identified as sufficient for specific spatiotemporal transcriptional activation of *MtCP6* in nodules. The functionality of this *cis*-regulatory sequence, thereafter named Nodule Senescence (NS), was validated by both gain- and loss-of-function approaches.
- ERF091, an AP2/ERF family transcription factor, was identified in a yeast one-hybrid (Y1H) screen as an NS-box interacting factor, and shown to mediate transcription activation of a NS-box:GUS reporter in transactivation assays in *Nicotiana benthamiana*.
- This work uncovered a new senescence-related nodule specific *cis-*regulatory region (NS-box) and provided evidence for the likely involvement of a stress-related ERF family member in the regulation of *MtCP6*, at the onset of nodule senescence.

## Introduction

Leguminous plants can offer important benefits for agriculture sustainability due to their ability to establish symbiotic associations with soil nitrogen-fixing bacteria of the Rhizobiaceae family (Yan & Bisseling, 2024). This symbiosis leads to the development of a *de novo* root organ, the root nodule, in which bacteria, differentiated in bacteroids, are able to reduce atmospheric dinitrogen to ammonia to the benefit of their host plant in exchange of photosynthetic carbohydrates (Ferguson *et al*., 2010; Syska *et al*., 2019). Nodules can be classified as either indeterminate or determinate type according to the presence or not of a persistent nodule meristem. Determinate nodules are spherical and lack a persistent meristem. Whereas, indeterminate nodules are elongated with a persistent apical meristem (zone I), which ensures continuous nodule growth, followed by sequential histological distinct zones reflecting different stages of rhizobia and plant cell differentiation. In the infection zone (zone II), rhizobia are released from infection threads into plant cells, where they are surrounded by a plant-derived peribacteroid membrane in a new organelle-like structure called symbiosome. Thereafter, bacteroids generally undergo a terminal differentiation process in interzone II–III to form nitrogen-fixing bacteroids that can reduce atmospheric nitrogen into ammonia via their nitrogenase enzyme (zone III).

The lifespan of nodules is time-limited and regulated by nodule senescence a process characterized by a decline of nodule nitrogen fixation capacity and a coordinated death of both bacteria and plant cells (Puppo *et al*., 2005; Van de Velde *et al*., 2006; Perez Guerra *et al*., 2010; Kazmierczak *et al*., 2020). While in determinate nodules senescence occurs from the center and extends to the periphery along with aging, in indeterminate nodules senescence is characterized by the development of a distinct nodule senescence zone (zone IV), and this process is extremely sensitive to abiotic stresses (Kazmierczak *et al*., 2020; Puppo *et al*., 2005).

A key feature of nodule senescence in legumes is the triggering of proteolytic activities for cellular degradation (Alesandrini *et al*., 2003; Puppo *et al*., 2005; Groten *et al*., 2006; Kazmierczak *et al*., 2020; Yang *et al*., 2020). Although protease activity can be regulated by peptide-based protease inhibitors during this process (Sharma & Gayen, 2021; Hellinger & Gruber, 2019), in the *Medicago truncatula* symbiosis, transcriptomic analyses provide evidence that this process is also tightly regulated at the transcriptional level (Van de Velde *et al*., 2006; Perez Guerra *et al*., 2010; Sauviac *et al*., 2022). Indeed, a set of cysteine protease encoding genes, *MtCP1* to *MtCP6,* have their expression strongly induced at the onset of nodule senescence in the *Medicago truncatula* - *Sinorhizobium meliloti* symbiosis. Further functional and expression characterization of *MtCP6* in *Medicago* has confirmed its importance as an early gene marker of both developmental and induced nodule senescence (Pierre et al., 2014).

Nodule development involves extensive transcription reprogramming controlled by major transcription factors (TF). Many of these TFs are crucial for regulating early stages of nodule development, rhizobial infection or bacteroid differentiation and nitrogen fixation (e.g. ERN1/2, CYCLOPS, NF-YA1, NIN, RSD, FUN, …) (e.g. Andriankaja et al., 2007, Middleton et al., 2007; Cerri et al., 2016, 2017; Sing, 2014; Hirsh et al., 2009; Vernié et al., 2015; Laporte et al., 2014; Liu et al., 2019; Sinharoy et al., 2013; Lin et al., 2024). However, only a limited number of TFs have been implicated in nodule senescence.

Notably, the MtbHLH2 transcription factor, which negatively regulate the cysteine protease encoding gene *MtCP77*, is down regulated during nodule senescence (Deng *et al*., 2019) most likely to allow *MtCP77* nodule senescence expression. Positive regulators such as NAC transcription factors play significant roles in nodule senescence (Yu et al, 2023; Wang et al, 2023a; Wang et al., 2023b; Xiao et al., 2024; de Zelicourt et al., 2012). In *M. truncatula*, *MtNAC969* is differentially regulated by salt stress and nodule senescence, and its downregulation via RNAi led to a premature nodule senescence phenotype associated with massive expression of various cysteine protease genes (de Zelicourt et al., 2012). In *Lotus japonicus*, after nitrate treatment, the *LjNAC094* gene was discovered as a regulator of nodule senescence and of senescence-associated genes (SAGs) (Wang *et al*. 2023a). In *Glycine max* several NACs were also shown to activate the expression of senescence-associated genes, including cysteine protease genes, during both developmental and nitrate-induced nodule senescence processes (Wang et al., 2023b; Xiao et al., 2024). More recently, new ERF transcription factors of group III, named *GmENS1* and *GmENS2 (*Ethylene-responsive transcription factors required for Nodule Senescence), were involved in the transcriptional regulation of NAC genes *GmNAC018*, *GmNAC030* and *GmNAC039* in soybean during nodule senescence (Xiao et al., 2024). Four NAC transcription factors (SNAPs) act as central regulatory hubs of nodule senescence induced by nitrate. SNAPs activate the expression of various transcription factors, among them seven ERFs, during nodule senescence (Wang et al., 2023b). These studies also led to the identification of *cis*-binding sites for nodule senescence-related transcription factors in soybean (Wang et al., 2023b; Xiao et al., 2024), although these *cis*-elements were not specifically validated during nodule senescence.

To gain insights into senescence-related promoter regulatory sequences, in this study in this study, we performed a detailed functional promoter analysis of *MtCP6,* the earliest marker of nodule senescence in *M. truncatula.* Serial analyses of *MtCP6* promoter deletions fused to the *GUS* reporter during the onset of developmental and nitrate-induced nodule senescence revealed that a *MtCP6* −242 bp proximal promoter upstream of the transcription start site (TSS) is sufficient to drive nodule senescence-associated expression of *MtCP6* in *Medicago*. We further defined through gain- and loss-of-function approaches that a 67 base pair sequence, termed the Nodule Senescence (NS) box or NS-box, is required and sufficient to drive tissue-specific expression of *MtCP6* during both developmental and nitrate induced nodule senescence. Finally, the stress-related ERF091 transcription factor was identified as an NS-box-interacting factor in a yeast-one-hybrid screen and shown in *Nicotiana benthamiana* assays to mediate NS-box transcriptional activation. Taken together, these results provide a novel senescence related cis-acting element and suggest the involvement of a stress-related ERF regulator in the transcriptional regulation of *MtCP6* during early nodule senescence in *M. truncatula*.

## Materials and Methods

### Biological material, root transformation and growth conditions

*Escherichia coli* DH5α was cultivated in LB Broth at 37℃ with appropriate antibiotics. *Sinorhizobium meliloti* 2011 was grown in LB (LB Broth, added with 2.5mM CaCl_2_ and 2.5 mM MgSO_4_) with 100µg/mL streptomycin and 10µg/mL tetracycline at 30℃. *Agrobacterium rhizogenes Arqua1* was grown in TY medium (5g/L bacto-tryptone, 3g/L yeast extract, 6mM CaCl_2_) with 100μg/mL streptomycin at 28 ℃(Quandt, 1993). *Agrobacterium tumefaciens* GV3101 and GV3103 strains were grown in LB Broth under antibiotic selection at 28°C (50μg/mL rifampicin, 15μg/mL gentamicin). YM4271 yeast strain was used in the Yeast-One-Hybrid screen (Matchmaker one-hybrid system, Clontech).

*M. truncatula* Jemalong A17 seeds were scarified, and surface-sterilized and germinated as described by Boisson-Dernier *et al*. (Boisson-Dernier *et al*., 2001). Germinated seedlings were stabbed at hypocotyl with *A. rhizogenes* Arqua1 suspension. The plantlets were thereafter transferred into a substrate mixture of perlite and sand (3:1) with a basic nitrogen supply of 1 mM NH_4_NO_3_. Plants were grown at 20℃ for one week for optimal transformation and then grown at 23℃ for 2 weeks. Transgenic roots of composite plants were selected by cutting off the non-green-fluorescent roots (GFP) under a Leica MZ FLIII fluorescence stereomicroscope (Leica). After one-week adaptation following the selective cutting, transgenic composite plants were inoculated with 10 mL of *S. meliloti* 2011 pXLGD4 (Grefen *et al., 2010*) at optical density of OD_600nm_= 0.1. Nodules were harvested at 4 week-post-inoculation (wpi). Nitrogen-treated nodules were from plants that were treated with 10 ml KNO_3_ (10mM) for 2 days before harvesting at 4 wpi.

### Plasmid constructions

Thirteen promoter fragments (−1,720bp, −1,467bp, −1,278bp, −1,088bp, −599bp, −511bp, −356bp, −303bp, −273bp, −242bp, −175bp, −141bp, and −80bp to the TSS) were amplified and cloned into the entry vector *pDONR*™ P4-P1r (Invitrogen) (Table S1). Empty entry vector was built by recombination of attB4::empty cassette::attB1 into *pDONR*™ P4-P1r. Expression vector was constructed by multi-site Gateway recombination of promoter entry vectors with *pENTR*-*GUS* and *pENTL2L3*-*T35S* into the destination vector *pKm43GWD*-*RolD*:*eGFP* (Cam et al., 2012).

For gain of function experiments, seamless tetramers of NS region (−242bp to −175bp) was synthesized and cloned into *pUC53-Kan* (Genewiz corporation). Monomer or Tetramers were then inserted into EcoRI/HindIII sites upstream of a minimal *CaMV35S* promoter (47bp) in a binary vector pLP100 (Szabados *et al*., 1995) according to Andriankaja *et al*. (2007) (Table S2). Binary constructs were then introduced into *A. rhizogenes* Arqua1 by electro-transformation (MicroPulser, Bio-Rad). Block deletion of the NS region was generated with complementary primers (Table S1) from *pDONR*™ P4-P1r-*ProCP6* (−1,720bp). Then the deleted promoters (ΔNS) was recombined into *pKm43GWD*-*RolD*:*eGFP* as described in promoter deletion construction (Table S2).

For transactivation experiment, BP reactions were performed according to the manufacturer’s instructions (Invitrogen) between ERF091 and ERF092 pGAD clones and the respective donor plasmid pDONR207 (Invitrogen Life Sciences). LR recombination reactions between pDONR207 constructs and destination vectors *pAmPAT-P35S*-3HA was done to generate the respective *pAmPATP-35S*-3HA-ERF091 and *pAmPAT-P35S*-3HA-ERF092 fusion constructs (Table S2).

### Histological analyses and microscope observations

Nodules with attached root fragments were harvested at 4 wpi, and at 4 wpi with a 2d-nitrate treatment. Samples were stained with X-Gluc (5-bromo-4-chloro-3-indolyl-β-D glucuronide, cyclohexylammonium salt; Euromedex) with a modified protocol from Jefferson et al. (Jefferson, 1987). Nodules were incubated in GUS staining solution at 37°C overnight. Alternatively, short-time staining (4h at 37°C) were applied to better analyze staining intensity of promoter deletions of less than −303 bp. Stained nodule images were taken with a Leica MZFLIII stereomicroscope (Leica). Stained nodules were thereafter embedded in 6% (w/v) agarose and sectioned (70 µm) with a HM650V vibratome (Thermo Fisher Scientific). Nodule sections were observed under dark field using the Axioplan 2 microscope (Carl Zeiss). From all histological experiments, at least 200 nodules were analyzed from more than 15 independent plants of three biological replicates.

### Yeast One-Hybrid Screening

A Yeast-One-Hybrid (Y1H) screen was performed using yeast strains carrying the 4x NS-box construct. The generation of the strain and the Y1H screen were done according to established methods (Andriankaja *et al*., 2007). The 4x NS-box was cloned into the pHISi vector to create the 4xNS-box-HIS3 construct, which was then integrated into the yeast genome. The Y1H screen was performed using 20 µg of a nodule AD fusion cDNA library generated from 4- and 19-days-old nodules (Baudin *et al*., 2015) and candidate colonies were selected on SD His^-^ Leu^-^ selective medium supplemented with a 10 mM 3-amino-1,2,4-triazole (3-AT) after 4 days incubation at 28°C. These colonies were then re-spotted on the same selective and non-selective media, and positive candidates were sequenced and analyzed against genomic databases (Fig. S3). Plasmid DNAs from selected NS-box-interacting ERF candidates were extracted and used to transform yeast bait strains 4x NS-box (this study) and 4x NF-box (Andriankaja *et al*., 2007) to validate their interaction with the NS-box relative to the ERN1 TF control. Dilution series (DO_600nm_= 1, 0.1 and 0.01) of the yeast transformants were spotted on selective media SD Leu^-^, SD His^-^Leu^-^, and SD His^-^Leu^-^ + 15 mM 3-AT and their growth were then monitored for 3-4 days, for evaluating the specificity of the interaction towards the 4x NS-box or 4x NF-box regulatory sequences.

### Transient Expression *in N. benthamiana* Leaves

*A. tumefaciens* GV3101 containing pLP100 construct harboring the gain-of-function 1X NS-box::GUS fusion and GV3103 strains containing pAmPAT-35S binary vectors harboring 3-HA tagged ERF91 protein were infiltrated in *N. benthamiana* leaves as described previously (Andriankaja et al., 2007). As positive control for transactivation experiments, 4X NF-box::GUS fusion and the HA-ERN1 protein were used (Andriankaja et al., 2007, Table S2). Leaf discs were harvested 36 h post-infiltration for direct histochemical GUS assays or frozen in liquid nitrogen prior to quantitative enzymatic GUS fluorometric assays or Western-blot analyses using Anti-HA-Peroxidase, High Affinity (3F10) monoclonal antibodies (Roche).

### Fluorometric GUS assays

20 mg of nodules were harvested and ground in liquid nitrogen. Total proteins were extracted with GUS buffer (50 mM sodium phosphate, pH 7.5, 10 mM 2-mercaptoethanol, 10 mM Na_2_EDTA, 0.1% Triton X-100, and 0.1% sodium lauryl-sarcosine). Quantitative GUS fluorimetric activities were measured using 10 µg of total protein extracts with 4-Methylumbelliferyl-β-D-glucuronide hydrate, (MUG) as substrate (Biosynth M-5700), as previously described by (Boisson-Dernier *et al*., 2005). Total protein from infiltrated *N. benthamiana* leaves disc were extracted, and GUS activities were measured using 1µg of total protein extract (Andriankaja *et al*., 2007) and 1 mM of the MUG substrate. GUS activities were measured using a FLUOstar Omega 96 microplate reader (BMG LABTECH). Standard curves were prepared with a range of increasing concentrations of 4-methylumbelliferone (4-MU) (Sigma-Aldrich).

### In silico analysis

Analysis was conducted with MEME algorithm (p-value>0.05) on MEME Suite web server (http://meme-suite.org<u>; Bailey et al., 2009;</u> Timothy et al., 1994) with the following parameters: maximum number of motifs (3), minimum motif width (6), maximum motif width (50), minimum sites per motif (2), maximum sites per motif (600).

### Statistic analysis

Our data, from Fig. 1 to Fig. 3, were reported as mean ± standard errors. The significance of the results was assessed using the ANOVA one-way with post-hoc Tukey HSD parametric test.

**Figure 1:**
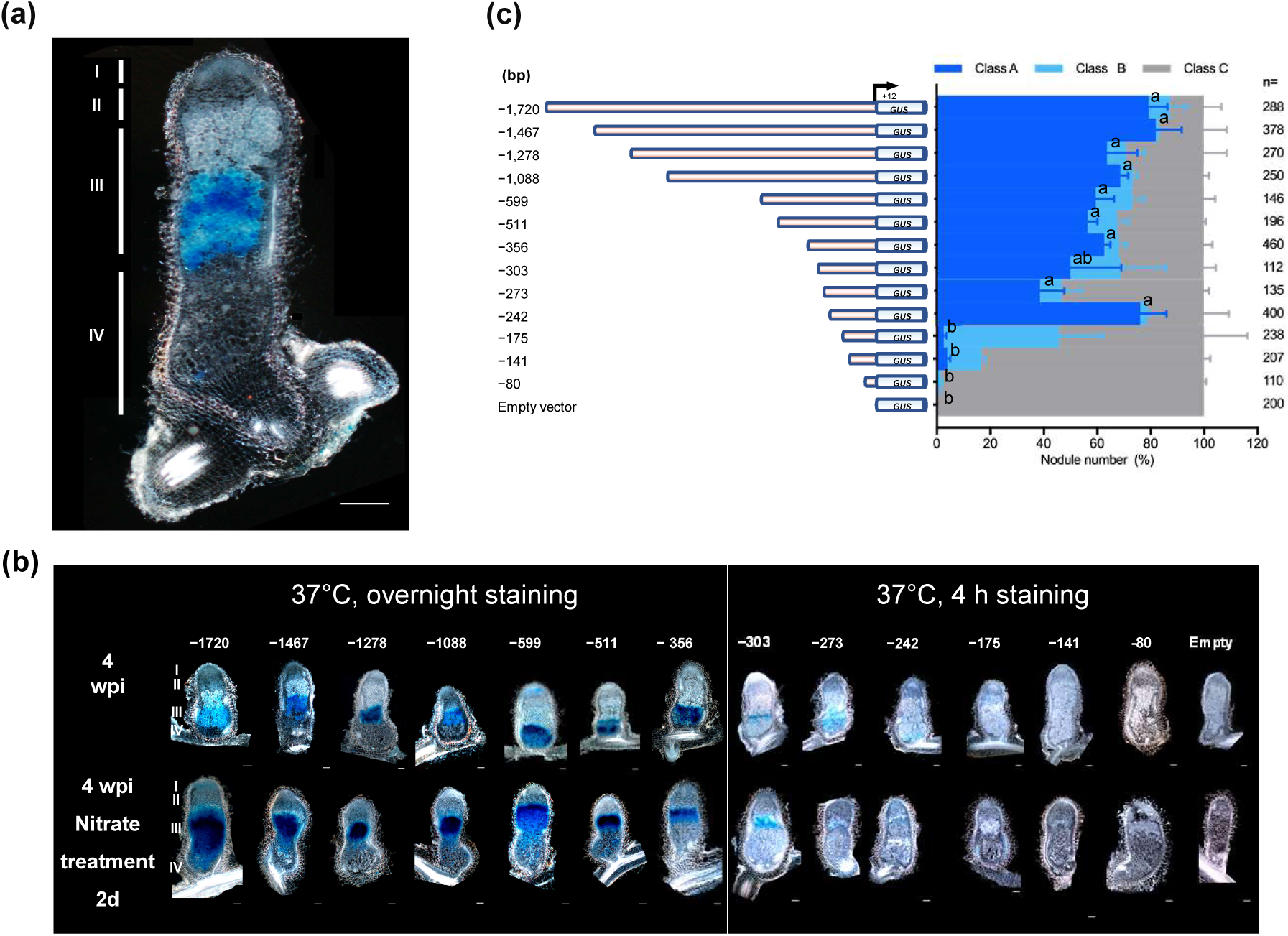
Schematic graph of *MtCP6* 5’-promoter deletion constructs, and corresponding GUS expression spatial specificity analysis. a. A representative image shows tissue-specific GUS activity driven by the 1,720 kb *MtCP6* promoter in the transition zone between zone III-IV of a 70 µm section of a nodule harvested 6 wpi (week-post-inoculation) from a *M. truncatula* transgenic root inoculated with *S. meliloti* 2011. **b.** Histochemical localization of GUS activity of *MtCP6* truncated promoters in nodule 70 µm sections at 4 wpi, 2d after nitrate treatment. Nodule sections were stained overnight (for −1,720, −1,467, −1,278, −1,088, −599, −511, and −356bp deletions), or during 4h (for −303, −273, −242, −175, −141, −80 bp deletions and empty vector). Promoter positions are indicated in bp relative to the TSS. Scale bars = 100 µm. **c.** Schematic depiction of progressive 5’deletions of *MtCP6* promoter constructs and empty vector. Spatial *GUS* expression pattern is shown in percentages and classed in specific expression in root nodule zone III-IV (blue, Class A), unspecific expression in other nodule zones (teal blue, Class B), and non-detectable (GUS-, grey, Class C). Total number of nodules analyzed is indicated on the right (n=110 to 460). GUS-stained nodules were harvested in more than three independent experiments. ANOVA one-way with post-hoc Tukey HSD parametric test was applied to the values (P < 0.05; n ≥ 110). Statistical differences among means are indicated by different letters.

The statistical analyses in Fig. 5 were performed using the R software (http://r-project.org). The data were first evaluated for normality using the Shapiro-Wilk test and the homogeneity of variances using the Fisher and Bartlett tests. The data in Fig. 5 showed a normal distribution but heterogeneity of variance. They were therefore analysed using non-parametric statistical tests, the Kruskal-Wallis test in Fig. 5A (Chisq=24.84355, p < 0.001) and 5B (Chisq=40.92735, p < 0.001) and the Mann-Whitney test in Fig. 5D (W=313, p < 0.001).

## RESULTS

### The truncated **−**242 bp *MtCP6* promoter region is sufficient to confer specific spatiotemporal *GUS* expression under both developmental and nitrate-induced nodule senescence

A −1720 bp *pMtCP6:GUS* fusion exhibit tissue specific expression in the nodule interface zone III-IV(ref) (Fig. 1a). In order to identify *cis*-regulatory elements responsible for this regulation, fifteen 5’ progressive deletions of the *MtCP6* promoter, ranging from −1,720 bp to −80 bp upstream of the Transcriptional Starting Site (TSS) were generated. These sequences fused to *GUS* gene were introduced in *M. truncatula* by *Agrobacterium rhizogenes* transformation and GUS staining was performed on transgenic roots and nodules harvested at 4 wpi (Fig. 1a, 1b). Nodules were classified according to relative GUS staining patterns in individual nodules. Class A nodules are those exhibiting specific GUS staining at the interzone III-IV (Fig. 1a). Unspecific expression in other nodule zones (Class B), and non-detectable (GUS-, Class C). This expression pattern was present in 87% of nodules for the full-promoter construct (−1,720 bp). Promoter deletions ranging from −1,467 bp to −242 bp (Fig. S1), showed similar interzone III-IV-specific expression profiles in 40-80 % of nodules and with no significant difference to the full-length promoter

Further 5’ deletions down to −211bp, drastically reduced the number of Class A-GUS stained nodules to 9 % (Fig. S1). Non-specific GUS staining predominantly appeared with shorter promoter deletions below −242 bp. At −175 bp, only 6 % of nodules presented a correct tissue specificity instead of more than 91% for −1,720 bp (Fig. 1c, Fig. S1). Our results suggest that *cis*-elements responsible for *MtCP6* transcription induction under developmental senescence are localized between −242 bp and −175 bp positions.

To identify *cis*-elements responsive to nitrate, nodules were collected two days after nitrate treatment. Under nitrogen treatment, GUS staining for the −1,720 bp *MtCP6* promoter construct expanded from proximal to distal nodule regions, indicative of an accelerated developmental senescence (Fig. 1b). A reduction in GUS activity was also observed for constructs containing promoter fragments at −242 bp and −175 bp. In conclusion, a promoter region spanning to −242 bp appeared sufficient to drive *MtCP6* gene transcription during both developmental and nitrate-induced nodule senescence. Thus, it is likely that senescence-associated *cis*-regulatory elements are localized within the −242 bp to −175 bp region of *MtCP6* promoter, termed as the “nodule senescence (NS) box” or NS-box (Fig. S2a).

### The NS-box is sufficient to endow nodule senescence-related expression patterns to the *GUS* reporter gene

Our findings suggest that the NS-box, consisting of a 67 bp sequence lying in between −242 bp and −175 bp positions of the *MtCP6* promoter, is sufficient to regulate the specific *MtCP6* spatiotemporal expression pattern associated with nodule senescence. To evaluate whether this sequence is sufficient for conferring this senescence-related expression pattern, we conducted gain-of-function experiments using a tetramer of the NS-box (4x NS). This tetramer was fused to a minimal CaMV 35S promoter (*Pmin35S*) and the GUS reporter gene within the pLP100 binary vector (Andriankaja et al., 2007; Fig. 2a). The expression pattern of the resulting gain-of-function construct was analyzed in transgenic root nodules at 4 wpi, with or without nitrate application. Representative images of transgenic nodules expressing the chimeric gene constructs are presented in Fig. 2b. GUS activity was observed in the proximal part of the nodule (Class A), when the GUS reporter gene was under the control of the NS tetramer (Fig. 2b, 2c). Under nitrate treatment, GUS activity was observed in 20% of nodules expressing the NS reporter, and all GUS-positive nodules displayed specific GUS staining localization (Fig. 2c). In contrast, no GUS staining was observed in nodules expressing the empty vector containing the *GUS* gene under the control of the minimal *CaMV* 35S promoter, *Pmin35S*. To validate this, GUS activity was quantified using fluorimetric GUS assays in isolated nodules expressing the NS tetramer or control reporters (Fig. 2d). A significant 6.9-fold increase in GUS activity (124±13 pmol MU /min/mg protein) levels was mediated by the 4xNS reporter in nodules compared to the *Pmin35S* control construct, whose basal activity were around 18±2 and 20±2 pmol MU/min/mg protein at 4 wpi (Fig. 2d). Upon nitrate treatment (4 wpi nitrate), GUS activity driven by the 4xNS reporter significantly increased over 39.3-fold to 786±110 pmol MU/min/mg protein compared to the *Pmin35S* control. Together, these data demonstrate that the tetramer of the NS-box fused to a minimal *CaMV 35S* promoter provides the tissue-specific senescence-related transcriptional regulation in nodules. This supports the hypothesis that the NS-box contains *cis*-regulatory elements mediating *MtCP6* transcription regulation during nodule developmental and nitrate-induced senescence. However, since the global expression level conferred by the 4xNS reporter is low compared to the full-length native promoter, it is likely that other *cis*-elements not included in the NS region are required for full *MtCP6* promoter activity.

**Figure 2:**
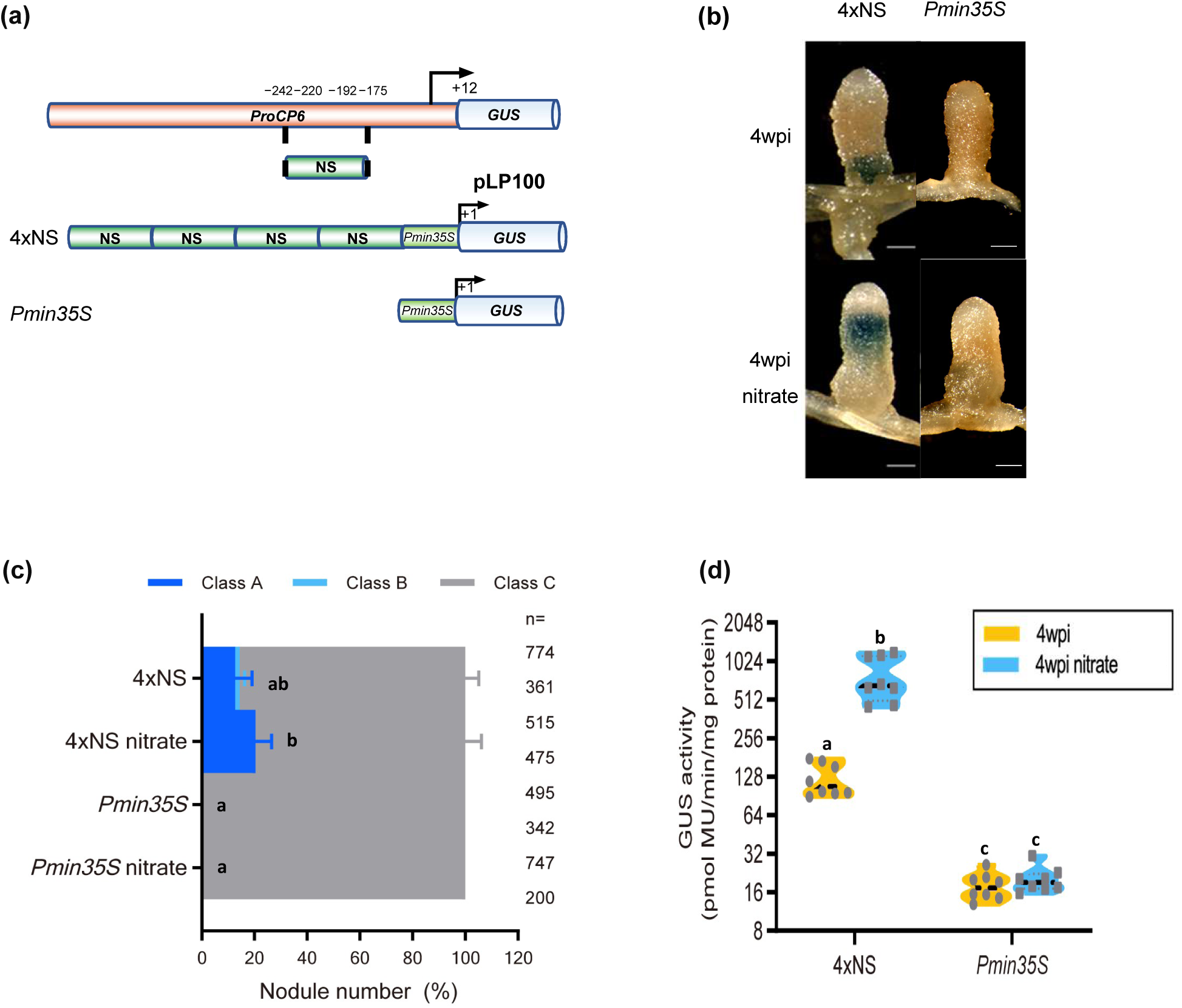
Gain of function analysis of *MtCP6 cis*-regulatory elements. **a.** Schematic representation of gain of function constructs. NS-box (Nodule Senescence, 67 bp) correspond to the promoter region from −242 to −175bp. Tetramers of NS (4x NS) was seamlessly synthesized and integrated in front of a minimal *CaMV 35S* promoter (*Pmin35S*, 47 bp). **b.** The images show nodules at 4wpi after GUS staining for 4x NS and control *Pmin*35S constructs. Blue staining shows the expression of *GUS* reporter gene in the transition from nitrogen-fixing zone III to senescent zone IV. **c.** Graphical representation of the overall spatial *GUS* expression patterns driven by the synthetic promoters (in root nodule zone III-IV (blue, Class A), unspecific expression in other nodule zones (teal blue, Class B), and non-detectable GUS signal (GUS-, grey, Class C). The 4-week-old nodules were harvested or treated with 10mM KNO_3_ for 2d. The total number of nodules analyzed is indicated on the right. GUS-stained nodules were harvested in more than three independent experiments. ANOVA one-way with post-hoc Tukey HSD parametric test was applied to the values (P < 0.05; n ≥ 200). Statistical differences among means are indicated by different letters. **d.** Quantitative fluorometric analyses of GUS activity driven by synthetic promoters in nodules at 4wpi with (blue) or without nitrate treatment (yellow). Data points stand for independent tests from three biological replicates. ANOVA one-way with post-hoc Tukey HSD parametric test was applied to the values (P < 0.05; n = 8). Statistical differences among means are indicated by different letters. Scale bar = 100 µm.

### Loss of the NS-box impairs *MtCP6* promoter transcriptional activity in nodules

Previous gain-of-function studies have suggested that *MtCP6* regulation during nodule senescence involves the NS-box and likely other *cis*-regulatory elements. To assess the importance of the NS-box regulatory region in the transcriptional activity of the −1,720 bp promoter, a loss-of-function analysis was conducted using a ΔNS deletion construct (Fig. 3a). Qualitative analysis of GUS activity revealed that transgenic nodules carrying the ΔNS construct still exhibited GUS activity in different nodules compared to the full promoter (Fig. 3b). However, quantitative analysis through fluorimetric assays showed a significant 6.1-fold decrease in GUS activity levels in ΔNS nodules (1,879±267 pmol MU/min/mg protein) compared to the −1,720 bp promoter construct (11,544±965 pmol MU/min/mg protein) (Fig. 3d). Although deletion of the NS region does not completely abolish the expression of *MtCP6* during nodule senescence, it drastically reduces its relative expression level. This suggests that *cis*-regulatory motifs in the NS-region and other promoter regions of *MtCP6* are required for its full expression during nodule senescence. Indeed, several motifs are found to be repeated along *MtCP6* promoter and might act together to amplify the expression of *MtCP6* at the onset of nodule senescence (Fig. S2b).

**Figure 3:**
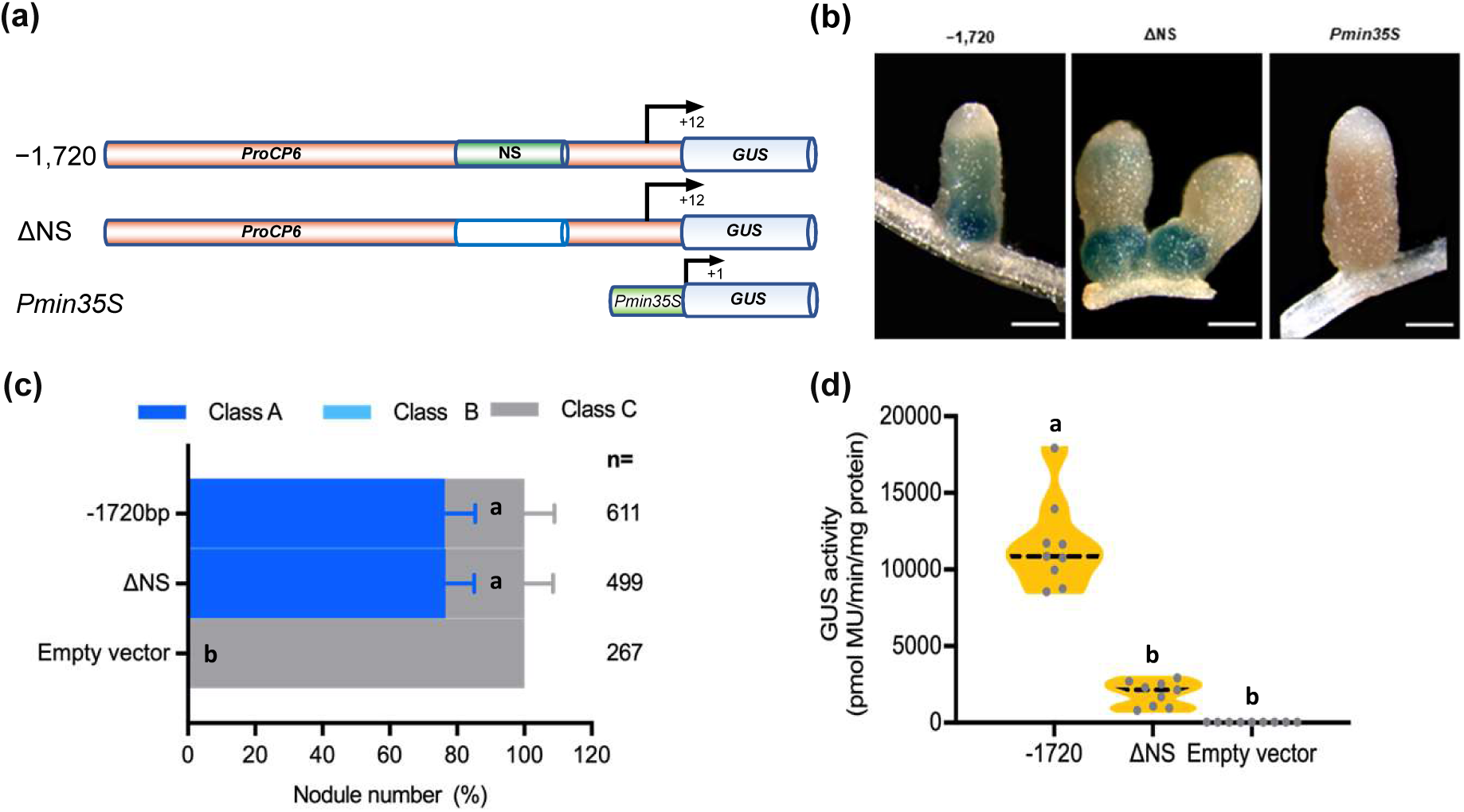
Loss of function of *cis*-regulatory elements on *MtCP6* promoter. **a.** Schematic representation of ΔNS deletion into the *MtCP6* promoter. The NS-box was deleted in the ΔNS construct from the *MtCP6* (−1,720 bp) promoter construct and then fused to the *GUS* reporter. **b.** Representative nodule images after GUS staining from the reporter constructs of *MtCP6* (−1,720 bp) promoter, ΔNS, and control *Pmin35S* constructs, respectively. **c.** Graphical representation showing respective percentages of nodules class A to C for *MtCP6* (−1,720 bp), the ΔNS and Pmin35S (empty vector) promoters (blue, Class A; teal blue, Class B; and grey, Class C). **d.** Fluorometric assay of GUS activity was performed with nodules expressing *ProCP6* (−1,720 bp), ΔNS, and control *Pmin35S* constructs at 4wpi. Results are obtained from nine independent experimental data points from three biological replicates. ANOVA one-way with post-hoc Tukey HSD parametric test was applied to the values (P < 0.05; n = 9). Statistical differences among means are indicated by different letters. Scale bar = 100µm.

### Identification of NS-box ERF-interacting transcription factors

To evaluate if DNA-binding protein factors can interact with the NS-box, a yeast-one-hybrid (Y1H) screen was performed using a tetramer of the NS-box (4xNS) as a bait. A *M. truncatula* nodule cDNA library (Baudin *et al*., 2015) was screened using a yeast strain YM4271 containing the HISTIDINE3 (HIS3) gene under the control of 4xNS fused to the HIS3 minimal promoter (Andriankaja et al., 2007). The bait strain, which is unable to grow under selective conditions (without leucine and histidine and supplemented with 5 mM 3-aminotriazole [3-AT]), allows efficient screening of the cDNA library for NS-binding TFs that can bypass this growth inhibition (Fig. S3). This screen led to the identification of seven positive NS-box interacting factors, all members of the ERF transcription factor family. Most positive clones (five out of seven) corresponded to two closely-related ERF091 (3/5 clones) and ERF092 factors (2/5 clones), belonging to group IX ERFs involved in plant defense and hormone signaling (Fig. S3d) (Middleton et al., 2007; Shu et al., 2018; Wu et al., 2022; Xie at al., 2019). The two remaining clones corresponded to ERF073 and ERF069, belonging to abiotic stress-related ERF VIII and anaerobic-related VII ERF factors, respectively (Fig. S3d). To confirm the specificity of the interactions, yeast transformation experiments were performed with isolated plasmids of the ERF candidates. Focus was on group IX ERF factors as they represented most of the positive NS-interacting factors identified in the Y1H screen study. As a control, another ERF transcription factor, ERN1, which was previously shown to interact with the NF-box *cis*-regulatory sequence in both Y1H and ChIP-PCR assays was used (Andriankaja et al., 2007; Cerri et al., 2016). Transformation of the yeast 4xNS tetramer bait strain with ERF091 or ERF092 expressing plasmids led to strong growth in selective conditions, indicative of ERF091/92 4xNS-bait interaction, compared to the ERN1control (Fig. 4a). Conversely, ERF091 or ERF092 were unable to interact with the NF-box, which was instead strongly recognized by ERN1, as previously shown (Andriankaja, 2007) (Fig. 4b). These results show the specific interaction of ERF091 and ERF092 with the NS-box regulatory sequence.

**Figure 4:**
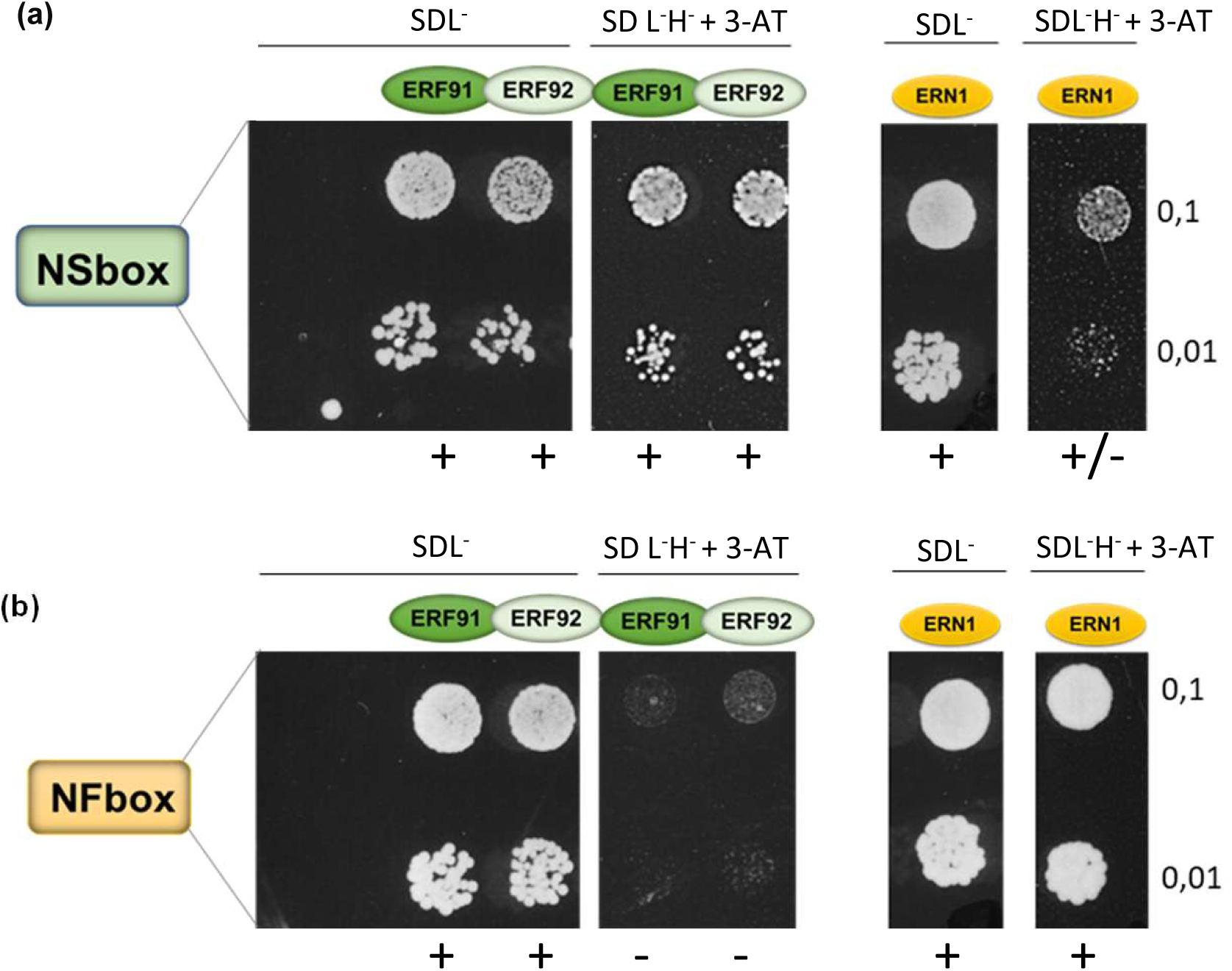
ERF091 and closely-related ERF092 interacts with the NS-box in Yeast-One-Hybrid (Y1H) assays. **a-b**. YM4271 yeast reporter strains carrying tetramers of NS-box (a) or NF-box (b) (Andriankaja et al., 2007) were transformed with plasmids expressing the ERF transcription factors ERF091, ERF092 or ERN1. ERN1, an interactor of the NF-box (Andriankaja et al., 2007, Cerri et al., 2016). Crossed yeast transformation studies of NS-box or NF-box yeast strains with the different factors were used to evaluate the relative specificity of ERF091 or ERF092 towards the NS-box. Yeast growth was examined in non-selective SD Leucine^-^ (SDL^-^) or selective conditions SD Histidine^-^ Leucine^-^ supplemented with 15 mM of 3-AT (SD H^-^ L^-^ + 3-AT). **(a)** The YM4271 NS-box tetramer strain transformed with ERF091 or ERF092 ERF constructs exhibited strong growth in selective SD H^-^ L^-^ + 3-AT conditions compared with the residual growth of the control NS strain transformed with ERN1. **(b)** The NF-box strain transformed with ERN1 showed expected strong growth in selective conditions while the same strain transformed with ERF091/092 plasmids did not show any significant growth. Representative images were taken 4 days after spot inoculation of yeast strains (at OD 0.1 and OD 0.01) transformed with pGAD plasmids expressing respective ERF factors (ERF091, ERF092 or ERN1). +, +/− and − indicate growth, residual growth or no significant growth, respectively. Non-transformed yeast control is marked as (−).

To determine whether these ERF IX regulators can regulate NS transcription in plant cells, transactivation experiments were performed in *Nicotiana benthamiana*. Since the tetramer 4xNS-box-GUS reporter showed background expression in this system, a monomer NS-box GUS reporter (NS-Box-GUS) was constructed for transactivation experiments in *N. benthamiana*. *N. benthamiana* leaves were infiltrated with *A. tumefaciens* strains carrying the NS-box-GUS reporter alone or with 3xHA-tagged ERF091 or ERN1 transcription factors expressed under the 35S promoter. As shown in Fig. 5a, ERF091 significantly activated the transcription of the NS-box-GUS reporter (median 276pmol MU/min/mg protein). Although ERN1 led to some crossed-activation of the NS-box-GUS reporter, this was relatively weaker (median 200pmol MU/min/mg protein) than the activation levels observed with ERF091. Conversely, only ERN1 led to a strong transactivation of the NF-box-GUS reporter compared to the residual transactivation by ERF091 (Fig 5b). Western blot analysis using anti HA antibodies confirmed that both ERF proteins were expressed in *N. benthamiana* leaf discs (Fig 5c). However, ERF091 was consistently produced at lower levels (∼1,7 x) than ERN1 in different experiments, as quantified using Imagelab software (BIO-RAD). To better represent relative GUS activity levels of ERF factors relative to their actual amounts in leaf discs, GUS activity levels (in Fig. 5a) were normalized against TF expression measured by protein band intensities (values obtained using Imagelab software. Normalized results show a 5.4x statistical difference that reinforce the conclusion of strong transcriptional activity of ERF091 towards the NS-box-GUS reporter compared to ERN1 (Figure 5d). In conclusion, ERF091 specifically interacts with and mediates transcriptional activation of the NS-box.

**Figure 5:**
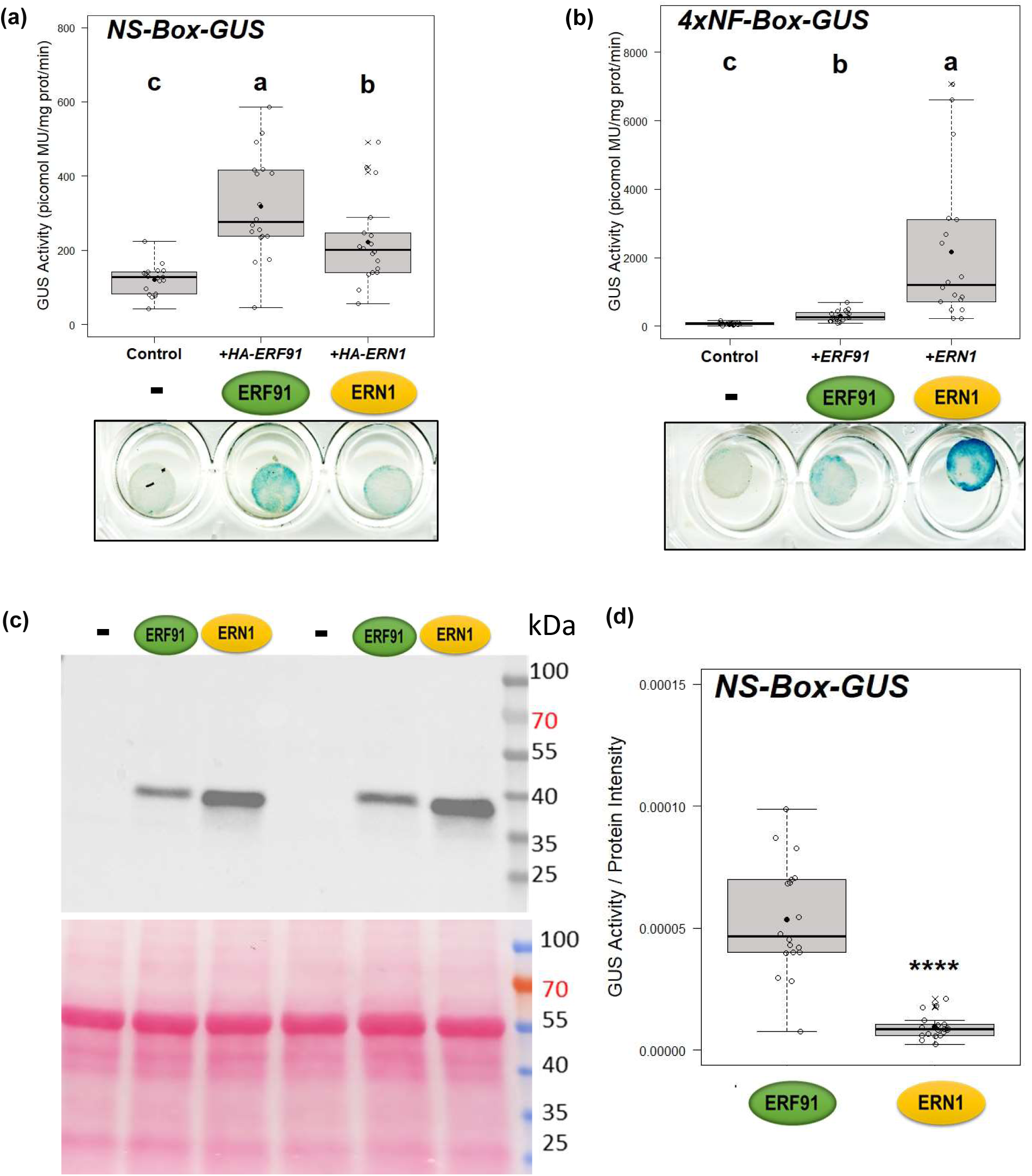
Transactivation of NS-box by ERF091 in *N. benthamiana.* **a**-**b**. Transactivation assays of p35Smin-GUS reporters fused to the NS-box-GUS (**a**) or to the NF-box tetramer (4X NF-box-GUS) (**b**) were performed in *Nicotiana benthamiana* epidermal leaf discs infiltrated with *A. tumefaciens* strains carrying promoter-GUS reporters only (−) or in the presence of respective ERF091 or ERN1 transcription factors, expressed under the CaMV 35S promoter. Fluorometric GUS assays were performed using 1 µg of total protein extracts from leaf discs. Box plots represent the distribution of values (indicated as open circles) of individual plants (n=18) from 3 independent experiments. Median (central line), mean (solid black circle) and outliers (cross) are indicated. Different letters indicate statistically significant difference (Kruskal-Wallis tests of the values were performed in R. P < 0.001). Representative images of histochemical GUS stained leaf discs are included. **c**. Relative protein expression levels of ERF91 and ERN1 were visualized in Western blot analysis (upper panel). Control of gel loading is shown after Ponceau S staining (lower panel). **d**. Box plots represent GUS activity levels of the NS-box-GUS reporter in the presence of ERF091 or ERN2 (shown in **a** and **b**) after normalization against relative protein band intensities of ERF091 and ERN1 (**c)**. A Mann-Whitney test was performed in R (asterisks indicate statistical difference; P < 0.001).

## DISCUSSION

To identify promoter sequences orchestrating spatiotemporal expression patterns at the onset of nodule senescence, a systematic serial deletion analysis of *MtCP6* promoter was conducted. Our starting point was the well-characterized −1,720 bp promoter of *MtCP6*, known to confer senescence and stress nitrate-induced expression in *Medicago* nodules (Perez Guerra et al., 2010; Pierre et al., 2014). In this study, we identified a sequence module spanning from −242 bp to −175 bp, termed the NS-box, sufficient to confer senescence-related expression in *Medicago* nodules when fused to a 35S min promoter (Fig. 2). Since the NS-box also responds to nodule nitrate treatment, this suggests that common *cis*-regulatory motifs within the NS-box mediate developmental and nitrate-induced senescence. Consistent with these results, ΔNS deletion in the 1,7 kb configuration resulted in a strong reduction of GUS activity levels (83% reduction). However, this did not completely abolished *MtCP6* promoter activity in nodules, suggesting that other motifs present beyond the NS region contribute to promoter activity. Further exploration is warranted to elucidate the role of these additional regulatory elements (Fig. S2).

In a Y1H screen using a NS-box tetramer, NS-box interacting factors have been identified, notably *MtERF091* and the closely-related *MtERF092,* members of the ERF class IX group, known for their roles in plant defense and hormone signaling (Wu et al., 2022). Previous ectopic expression of *MtERF091* in *Medicago* increased resistance to *Rhizoctonia solani* without an apparent impact on root nodulation (Anderson et al., 2010). In *L. japonicus*, *LjERF1*, the homolog of *MtERF091,* activates the expression of defense genes and positively impact nodule development in response to infection by *Mesorhizobium loti* (Asamizu et al., 2008). The discovery in this study that *MtERF091*, is able to interact and transcriptionally activate the senescence-related NS-box, suggests a possible involvement of this regulator in *MtCP6* gene expression during nodule senescence, and supports the hypothesis that nodule senescence might be associated with reactivation of plant defense responses. As nodules senesce, the weakening of the legume-rhizobia symbiosis may trigger a defense-like response. Ethylene, which is implicated in nodule senescence regulates the expression of the *MtERF091* homolog in *L. japonicus*, consistent with a possible involvement of *ERF091* in the ethylene-induced pathways promoting nodule senescence (Muller & Munne-Bosch, 2015; Shu et al., 2015; Phukan et al., 2017; Shu et al., 2018; Xie et al., 2019).

Although MtERF091 binds to the NS-box and promotes its transcription activity, a typical GCC-box cis-motif (AGCCGCC, Ohme-Takagi and Shinshi, 1995), bound by ERF IX TFs of other plant species, was not found in the NS-box (Franco-Zorilla et al., 2014; Wu et al., 2022). However, it is possible that MtERF091 recognizes other secondary *cis*-motifs outside the canonical GCC motif, as has been shown for other ERF family members (Franco-Zorilla et al., 2014). In this context, two GCC-rich motifs located within and just above the NS-box (shown as teal blue rectangles in Fig. S2), could potentially contribute to MtERF091-mediated *MtP6* transcriptional regulation. Future mutagenesis of these sites and DNA binding studies should help determine their relative importance. Consistent with our findings, two ERF transcription factors have recently been reported as key regulators of nodule senescence in *Glycine max* (Xiao et al., 2024). These ERF regulators, termed *GmENS1* and *GmENS2*, also bind to GC-rich promoter regions, although the associated senescence-specific *cis*-regulatory motifs remain to be determined.

In conclusion, this study has identified a specific *cis*-regulatory module for nodule-senescence, the NS-box, which in a synthetic form represents a valuable marker for tracking the onset of nodule senescence in *M. truncatula*. MtERF091/92 ERF transcription factors, identified here as novel NS-box interacting and transactivation factors, thus represent new potential regulators of nodule senescence in *Medicago*. The dual regulatory role of NS-box in developmental and nitrate-induced senescence suggests potential signalling overlap between plant defence and nodule-aging regulated pathways. Future identification of *cis*-regulatory motifs recognized by ERF091/92 within the NS-box may help to elucidate regulatory networks controlled during developmental or aging-related nodule senescence and potential signalling interplay with plant immunity.

## ACKNOWLEDGEMENTS

LY is supported by a doctoral fellowship from the China Scholarship Council (CSC). This work was supported by the “Institut National de la Recherche Agronomique”, the “Université Côte d’Azur”, the French Government (National Research Agency, ANR) through the “STAYPINK” project reference # ANR-15-CE20-0005), the “Investments for the Future” LABEX SIGNALIFE: program reference # ANR-11-LABX-0028-01, IDEX UCAJedi ANR-15-IDEX-01 and by National Natural Science Foundation of China (No. 32300211). This work was supported by French National Research grants TULIP ANR-10-LABX-41 to LF and FdCN. We are largely indebted to the symbiosis team members for numerous nodule harvests.

## AUTHOR’S CONTRIBUTION

Conceived the project (EB, PF). Promoter functional experiments: design and data analysis by EB, PF, LY supervised by EB, PF; Y1H and transactivation: design and data analysis by LY, LF, FdCN, supervised by FdCN; Performed experiments: LY, LF; analyzed data: LY, LF, FDC-N, EB, PF; wrote the paper: LY, FDC-N, EB, PF.

## DATA AVAILABILITY

The datasets generated for this study are available on request to the corresponding author.

## SUPPLEMENTARY DATA

**Table S1: List of primers used in this study.**

**Table S2: List of strains, plasmids and plasmid constructs.**

**Figure S1: Overall spatial GUS expression pattern.**

Percentages for each class of spatial GUS expression pattern in nodule. Root nodule specific expression in zone III-IV (Class A), unspecific expression in other nodule zones (Class B), and non-detectable (Class C). Total number of nodules analyzed is indicated on the right (n=110 to 460). GUS-stained nodules are from more than three independent experiments.

**Figure S2: NS box enrichment in co-expressed *CPs* (1755 bp promoter fragments).**

**(a) Nodule Senescence-box (NS-box, −242∼−175bp)**

**(b)** Research of motif similarities over *MtCP6* promoter using as Bioinformatic tool MEME Suite web server (http://meme-suite.org). Three repeated MEME Motifs are found onto *MtCP6* promoter (Bailey et al., 2009; Timothy et al., 1994). 1. TGAGRACCAMT (5 repeats in red); 2. CRWCGGCC (2 repeats in blue) and 3. GGTGTACC (2 repeats in green). Overlapping NS-box is mark with a dark blue box. IUPAC nucleotide code (A: Adenine, C: Cytosine, G: Guanine, T: Thymine, R: A /G, Y: C/T, S: G/C, W: A /T, K: G/T, M: A/C, B: C/G/T, D: A/G/T, H: A/C/T, V: A/C/G, N: any base).

**Figure S3: Schematic illustration of the Y1H screen to explore the 4xNS-box interactive proteins.**

**a.** The synthetic 4X NS-box promoter was integrated into the YM4271 yeast strain and screened from of a nodule cDNA library. **b**. Clones are isolated as able to grow on the SD His^-^ Leu^-^ + 10mM 3-AT selective medium, thereafter classified in three types (a type, b type, and c type). **c**. The positive clones are confirmed by another round of growth on selective medium, and the deduced amino acids are corresponding to the AP2/ERF family proteins. **d**. List of the seven MtERFs found and associated to their ERF family groups.

