## Supplemental Tables and figures for "Identification of regulatory promoter sequences directing *MtCP6* transcription at the onset of nodule senescence *in Medicago truncatula*"

### Table S1: Primer sequence list

| Primers | Sequences (5’ – 3’) | References |
|---|---|---|
| **−1720 F (attB4)** | GGGGACAACTTTGTATAGAAAAGTTGTCCGCCATCTATCTTCATTCGTCA | This work |
| **−1467 F (attB4)** | GGGGACAACTTTGTATAGAAAAGTTGTCTGAGAGAGATGAATGGTACTAT | This work |
| **−1278 F (attB4)** | GGGGACAACTTTGTATAGAAAAGTTGTCATGAGATGAGATGAGATGAGATGA | This work |
| **−1088 F (attB4)** | GGGGACAACTTTGTATAGAAAAGTTGTCGATAGTGATTGAGGACCGCT | This work |
| **−599 F (attB4)** | GGGGACAACTTTGTATAGAAAAGTTGTCTGAAATAAAGAGCTATTTGTTG | This work |
| **−511 F (attB4)** | GGGGACAACTTTGTATAGAAAAGTTGTCTTGTAAAAAATAATCAAAATTATCTA | This work |
| **−356 F (attB4)** | GGGGACAACTTTGTATAGAAAAGTTGTCCCATAGAATTTGCCATCACAAT | This work |
| **−303 F (attB4)** | GGGGACAACTTTGTATAGAAAAGTTGTCCATCACTTAGTTACGTGAAGCA | This work |
| **−273 F (attB4)** | GGGGACAACTTTGTATAGAAAAGTTGTCACACGTTAGTTAACGTGAAACA | This work |
| **−242 F (attB4)** | GGGGACAACTTTGTATAGAAAAGTTGTCTTGATTGTGACATACTTTCTTT | This work |
| **−175 F (attB4)** | GGGGACAACTTTGTATAGAAAAGTTGTCGAAAATTCTCACATTCTCACAG | This work |
| **−141 F (attB4)** | GGGGACAACTTTGTATAGAAAAGTTGTCGATTCCTACATTGGGTCTCATT | This work |
| **−80 F (attB4)** | GGGGACAACTTTGTATAGAAAAGTTGTCTTAGCACAAATTATTAACATAATA | This work |
| **attB1R-ProCP6** | GGGGACTGCTTTTTTGTACAAACTTGGAAGAAGTGCTGTGCTTTTACCC | This work |
| **empty vecteur F (attB4-attB3)** | GGGGACAACTTTGTATAGAAAAGTTGTCCAAGTTTGTACAAAAAAGCAGTCCCC | This work |
| **empty vecteur R (attB4-attB3)** | GGGGACTGCTTTTTTGTACAAACTTGGACAACTTTTCTATACAAAGTTGTCCCC | This work |
| **M13F** | GTAAAACGACGGCCAG | This work |
| **M13R** | CAGGAAACAGCTATGAC | This work |
| **pLP100-MSC F** | TGCCACCTGACGTCTAAGAAAC | This work |
| **GUS R** | AAGACTTCGCGCTGATACC | This work |
| **ΔNS 5' R** | CTGTGAGAATGTGAGAATTTTCTAACTCAGTTTGTTTCACGTTAAC | This work |
| **ΔNS 3' F** | GTTAACGTGAAACAAACTGAGTTAGAAAATTCTCACATTCTCACAG | This work |
| **ΔCW F** | CTTTGTATAGAAAAGTTGTCTTGTACTTTCTTTGTAAGAAAAATGC | This work |
| **ΔCW R** | GCATTTTTCTTACAAAGAAAGTACAAGACAACTTTTCTATACAAAG | This work |
| **ΔDOF F** | GAAAAGTTGTCTTGATTGTGACATAGTAAGAAAAATGCGTTCACTTGACA | This work |
| **ΔDOF R** | TGTCAAGTGAACGCATTTTTCTTACTATGTCACAATCAAGACAACTTTTC | This work |
| **pHIS fw (LS) (Lisa F)** | TTCCCAGTCACGACGTTG | This work |
| **pHIS rev (LS) (Lisa R)** | ATATTCTTCGAAGAAATCAC | This work |
| **pGAD10-R1** | CAC AGT TGA AGT GAA CTT GC | This work |
| **pGADT7-For1** | AGT ACC CAT ACG ACG TAC CAG | This work |

#### Table S2: List of strains and Plasmids

| Strains/plasmids | Characteristics | References |
|---|---|---|
| **Strains** | | |
| ***A. rhizogenes* ARqua1** | *Sm*$^R$, derivative of R1000 strain, moderate virulence | (Quandt, 1993) |
| ***A. tumefaciens* GV3101** | *Rif*$^R$ *Gen*$^R$ | (Van Larebeke et al., 1974) |
| ***A. tumefaciens* GV3103** | *Sm*$^R$ *Rif*$^R$ *Gen*$^R$ | (Holsters et al., 1980) |
| ***E. coli* DH5α** | F⁻ ϕ80*lac*ZΔM15 Δ(*lac*ZYA-*arg*F)U169 *rec*A1 *end*A1 *hsd*R17($r_K^-$, $m_K^+$) *phoA* *sup*E44 λ⁻*thi*$^{-1}$ *gyr*A96 *rel*A1 | Invitrogen |
| ***S. cerevisiae* YM4271** | Yeast reporter strain *MATa, ura3–52, his3–200, ade2–101, lys2–801, leu2–3, 112, trp1–901, tyr1–501, gal4-Δ512, gal80-Δ 538, ade5::hisG* | Clontech |
| ***S. meliloti* 2011** | *Sm*$^R$ | (Rosenberg et al., 1981) |
| **Plasmids** | | |
| **pDONR207** | *Gen*$^R$, entry vector for Gateway cloning | Invritogen |
| **pKGWFS7** | *Sp*$^R$, destination vector for Gateway cloning (*GOI::GUS::GFP*) | Invritogen |
| **pDONR P4-P1R** | *Km*$^R$, 5' entry vector for three entry multisite Gateway cloning | Invritogen |
| **pENTR-GUS** | *Km*$^R$, gene entry vecteur contains *uidA* (*GUS*) gene | Invritogen |
| **pENTR-T35S** | *Km*$^R$, 3' entry vecteur contains the *35S* terminator | Invritogen |
| **pKM43-rolD::GFP** | *Sp*$^R$, destination vector for three entry multisite Gateway cloning (pKM43) | (Karimi et al., 2002) |
| **pLP100** | *Km*$^R$, –47bp *Camv35S* (*pmin35S::GUS*) | (Andriankaja et al., 2007) |
| **pHISi** | *Amp*$^R$, HIS3, reporter vector for Yeast-One-Hybrid | Clontech |
| **pGAD-HA** | *Amp*$^R$, *Leu*, prey vector for Yeast-One-Hybrid | Clontech |
| **pAmPATp35S-3HA** | *Amp*$^R$, *Cb*$^R$, destination vector with 3HA tag | (Andriankaja et al., 2007) |
| **Plasmid constructions** | | |
| **pKM43-1720** | pCP6 -1,720bp, Gateway construction plasmid pKM43 | This study |
| **pKM43-1467** | pCP6 -1,467bp, Gateway construction plasmid pKM43 | This study |
| **pKM43-1278** | pCP6 -1,278bp, Gateway construction plasmid pKM43 | This study |
| **pKM43-1088** | pCP6 -1,088bp, Gateway construction plasmid pKM43 | This study |
| **pKM43-599** | pCP6 -599bp, Gateway construction plasmid pKM43 | This study |
| **pKM43-511** | pCP6 -511bp, Gateway construction plasmid pKM43 | This study |
| **pKM43-356** | pCP6 -356bp, Gateway construction plasmid pKM43 | This study |
| **pKM43-303** | pCP6 -303bp, Gateway construction plasmid pKM43 | This study |
| **pKM43-242** | pCP6 -242bp, Gateway construction plasmid pKM43 | This study |
| **pKM43-175** | pCP6 -175bp, Gateway construction plasmid pKM43 | This study |
| **pKM43-141** | pCP6 -141bp, Gateway construction plasmid pKM43 | This study |
| **pKM43-80** | pCP6 -80bp, Gateway construction plasmid pKM43 | This study |
| **pKM43-0** | Empty construct, Gateway construction plasmid pKM43 | This study |
| **pKM43-ΔNS** | pKM43-ΔNS:GUS:T35S, Gateway construction plasmid pKM43 | This study |
| **pLP100-Pmin35S-GUS** | Empty control | This study |
| **4X NFbox::GUS** | 4X NFbox::GUS positive control | (Andriankaja et al., 2007) |
| **pLP100-4xNS** | 4xNS, binary vector | This study |
| **pLP100-4xNS1** | 4xNS1, binary vector | This study |
| **pLP100-4xNS2** | 4xNS2, binary vector | This study |
| **pLP100-1xNS** | 1xNS, binary vector | This study |
| **pHISi 4xNS** | yeast bait, 4xNS | This study |
| **pGAD-ERN1** | Positive control, ERF V in pGAD plasmid | (Andriankaja et al., 2007) |
| **pGAD-ER091** | ERF IX in pGAD plasmid | This study |
| **pGAD-ER092** | ERF IX in pGAD plasmid | This study |
| **pGAD-ER073** | ERF VII in pGAD plasmid | This study |
| **pGAD-ER069** | ERF VII in pGAD plasmid | This study |
| **pAmPAT-ERF91** | ERF IX in pAmPAT plasmid (pAmPATp35S-3HA-ERF091) | This study |
| **pAmPAT-ERF92** | ERF IX in pAmPAT plasmid (pAmPATp35S-3HA-ERF092) | This study |

**Fig. S1**

| ProCP6 | n= | Class A % | Class B % | Class C % | GUS+ (Class A+B) % | Tissue specificity (Class A/GUS+) % |
|---|---|---|---|---|---|---|
| **-1720** | 288 | 79 | 8 | 13 | 87 | 91 |
| **-1467** | 378 | 82 | 1 | 17 | 83 | 99 |
| **-1288** | 270 | 64 | 7 | 29 | 71 | 90 |
| **-1098** | 250 | 69 | 5 | 26 | 74 | 94 |
| **-599** | 146 | 59 | 14 | 27 | 73 | 81 |
| **-511** | 196 | 57 | 11 | 32 | 68 | 84 |
| **-356** | 460 | 63 | 6 | 32 | 68 | 92 |
| **-303** | 112 | 50 | 19 | 31 | 69 | 73 |
| **-273** | 135 | 39 | 8 | 53 | 47 | 83 |
| **-242** | 400 | 76 | 3 | 21 | 79 | 97 |
| **-211** | 239 | 9 | 1 | 90 | 10 | 90 |
| **-175** | 238 | 3 | 43 | 54 | 46 | 6 |
| **-141** | 207 | 4 | 13 | 83 | 17 | 24 |
| **-80** | 110 | 0 | 1 | 99 | 1 | 0 |
| **0** | 200 | 0 | 0 | 100 | 0 | 0 |

**Fig. S2**

**(a)** NS-Box

```
   -242                  -220                        -192             -175
5'-TTGATTGTGACATACTTTCTTTGTAAGAAAAATGCGTTCACTTGACAAAGAATTTGGCCGTTGAATAG-3'
3'-AACTAACACTGTATGAAAGAAACATTCTTTTTACGCAAGTGAACTGTTTCTTAAACCGGCAACTTATC-5'
```

**(b)**

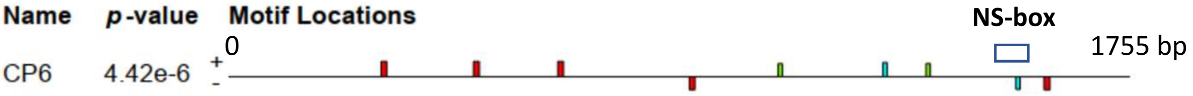


| Motif | Symbol | Motif Consensus |
|---|---|---|
| 1. | (red) | TGAGRACCAMT |
| 2. | (cyan) | CRWCGGCC |
| 3. | (green) | GGTGTACC |

1.
*E*-value: 8.9e+002 Site Count: 5 Width: 11

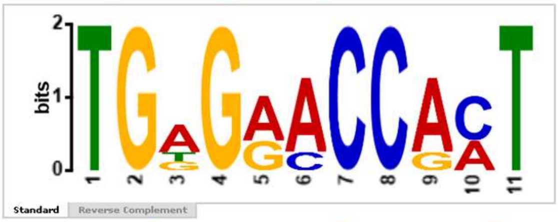


Log Likelihood Ratio: 66 Information Content: 17.2 Relative Entropy: 19 Bayes Threshold: 8.44294

| Name | Strand | Start | *p*-value | Sites |
|---|---|---|---|---|
| 1. CP6 | + | 642 | 2.39e-7 | TGATAGTGAT TGAGGACCGCT CCCAATGACA |
| 1. CP6 | + | 478 | 2.39e-7 | ATGGAGCATA TGAGAACCACT TGAGAGACTA |
| 1. CP6 | + | 298 | 3.07e-7 | TTAGAGCATA TGGGAACCACT TAAAGACTAA |
| 1. CP6 | - | 1589 | 1.85e-6 | ATTTTCTGAA TGAGACCCAAT GTAGGAATCT |
| 1. CP6 | - | 898 | 1.98e-6 | TATTCCTTTT TGTGGACCAAT GTGTTGTCAC |

2.
*E*-value: 6.9e+003 Site Count: 2 Width: 8

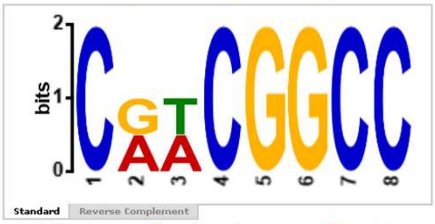


Log Likelihood Ratio: 26 Information Content: 14 Relative Entropy: 18.4 Bayes Threshold: 9.76984

| Name | Strand | Start | *p*-value | Sites |
|---|---|---|---|---|
| 1. CP6 | + | 1275 | 9.14e-7 | AATTTTATTC CGTCGGCC TTTTTGATAA |
| 1. CP6 | - | 1534 | 3.17e-6 | ATTTTCTATT CAACGGCC AAATTCTTTG |

3.
*E*-value: 7.9e+003 Site Count: 2 Width: 8

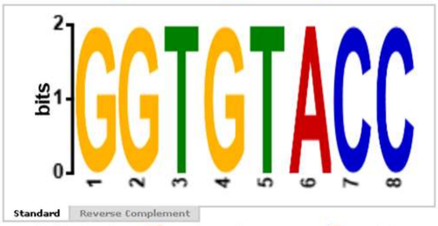


Log Likelihood Ratio: 26 Information Content: 16 Relative Entropy: 18.5 Bayes Threshold: 9.76984

| Name | Strand | Start | *p*-value | Sites |
|---|---|---|---|---|
| 1. CP6 | + | 1359 | 2.79e-6 | CAAATCTTTG GGTGTACC ATAGAATTTG |
| 1. CP6 | + | 1071 | 2.79e-6 | CACAAATTAA GGTGTACC GGTACTTTAA |

**Fig. S3** **Supplemental Figure: to illustrate the yeast one hybrid strategy**

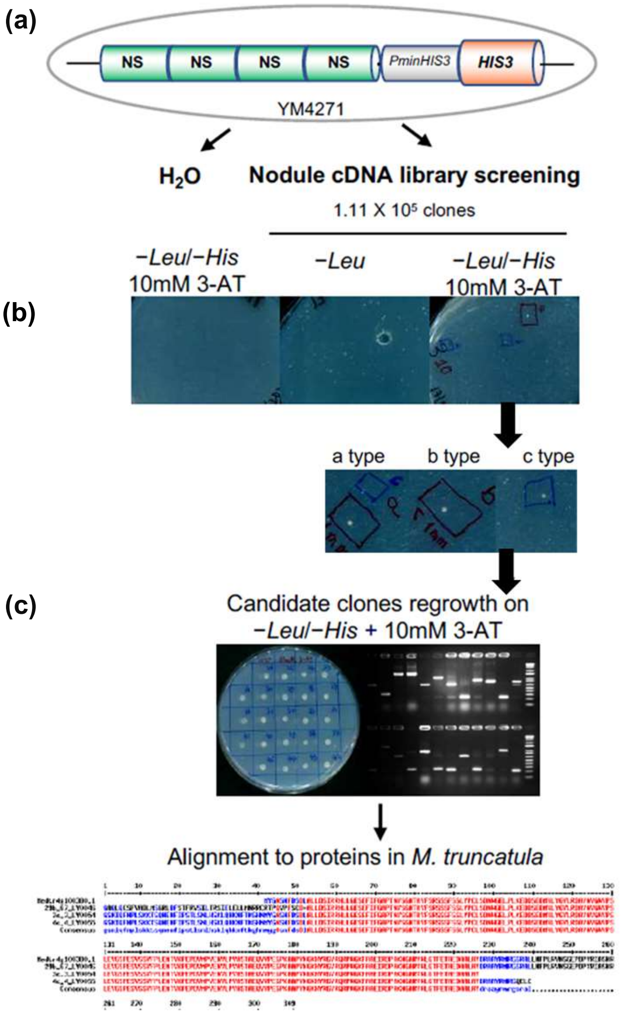


(d)

| Gene locus | Occ | Genome annotation | Name | Group |
|---|---|---|---|---|
| Medtr4g100380.1 | 3 | Ethylene-Responsive Factor | MtERF091 (MtERF1-1) | ERF IX |
| Medtr4g100420.1 | 2 | Ethylene-Responsive Factor | MtERF092 | ERF IX |
| Medtr2g435590.1 | 1 | Ethylene-Responsive Factor | MtERF069 (MtERFB2.3) | ERF VII |
| Medtr4g078710.1 | 1 | Ethylene-Responsive Factor | MtERF073 | ERF VIII |
